# Findings of a retrospective, controlled cohort study of the impact of a change in Nature journals' editorial policy for life sciences research on the completeness of reporting study design and execution

**DOI:** 10.1101/187245

**Authors:** Malcolm Robert Macleod, The NPQIP Collaborative group

## Abstract

**Objective:** To determine whether a change in editorial policy, including the implementation of a checklist, has been associated with improved reporting of measures which might reduce the risk of bias.

**Methods:** The study protocol has been published at DOI: 10.1007/s11192-016-1964-8.

**Design:** Observational cohort study.

**Population:** Articles describing research in the life sciences published in Nature journals, submitted after May 1st 2013.

**Intervention:** Mandatory completion of a checklist at the point of manuscript revision.

**Comparators:** (1) Articles describing research in the life sciences published in Nature journals, submitted before May 2013; (2) Similar articles in other journals matched for date and topic.

**Primary Outcome:** Change in proportion of Nature publications describing in vivo research published before and after May 2013 reporting the Landis 4 items (randomisation, blinding, sample size calculation, exclusions).

We included 448 NPG papers (223 published before May 2013, 225 after) identified by an individual hired by NPG for this specific task, working to a standard procedure; and an independent investigator used Pubmed Related Citations to identify 448 non-NPG papers with a similar topic and date of publication in other journals; and then redacted all publications for time sensitive information and journal name. Redacted manuscripts were assessed by 2 trained reviewers against a 74 item checklist, with discrepancies resolved by a third.

**Results:** 394 NPG and 353 matching non-NPG publications described in vivo research. The number of NPG publications meeting all relevant Landis 4 criteria increased from 0/203 prior to May 2013 to 31/181 (16.4%) after (2-sample test for equality of proportions without continuity correction, X^2^ = 36.2, df = 1, p = 1.8 x 10-9). There was no change in the proportion of non‐ NPG publications meeting all relevant Landis 4 criteria (1/164 before, 1/189 after). There were more substantial improvements in the individual prevalences of reporting of randomisation, blinding, exclusions and sample size calculations for in vivo experiments, and less substantial improvements for in vitro experiments.

**Conclusions:** There was a substantial improvement in the reporting of risks of bias in in vivo research in NPG journals following a change in editorial policy, to a level that to our knowledge has not been previously observed. However, there remain opportunities for further improvement.

## Background

Few publications describing *in vivo* research report taking specific actions designed to reduce the risk that their findings are confounded by bias, and those that do not report such actions give inflated estimates of biological effects. Strategies and guidelines which might improve the quality of reports of *in vivo* research have been proposed, [1,2] and while these have been endorsed by a large number of journals there is evidence that this endorsement has not been matched by a substantial increase in the quality of published reports [3].

Poor replication of *in vitro* molecular and cellular biology studies has also been reported [4,5] and this has been attributed in part to poor descriptions of the experimental and analytical details.

In May 2013 Nature Journals announced a change in editorial policy which required authors of submissions in the life sciences to complete a checklist, at the time of manuscript acceptance, indicating whether or not they had taken certain measures which might reduce the risk of bias and to report key experimental and analytical details; and in their submission to detail where in the manuscript these issues were addressed [6].

The aim of this study was to determine whether the implementation of this checklist for submissions has been associated with improved reporting of measures that might reduce the risk of bias. To establish whether any observed change in quality was a simply a secular trend occurring across all journals we matched each included publication with a publication in a similar subject area published at around the same time by a different publisher.

## Methods

The methods are described in detail in the published study protocol [7], and the data analysis plan and analysis code were articulated prior to database lock and registered on the Open Science Framework (https://osf.io/mqet6/#). The complete study dataset including PMIDs (but not, for copyright reasons, the source pdfs) of included articles is available on Figshare (10.6084/m9.figshare.5375275).

In this observational cohort study we aimed to determine whether the implementation of a checklist for submissions has been associated with improved reporting of measures which might reduce the risk of bias. The study populations comprised (1) Published articles accepted for publication in Nature journals, which described research in the life sciences and which were submitted after May 1^st^ 2013, at which time the mandatory completion of a checklist at the stage of manuscript revision, was introduced. This checklist required authors to indicate where details relating to study design could be found in the manuscript at the point of manuscript revision. and before November 1^st^ 2014; (2) Published articles accepted for publication in Nature journals in the months preceding May 2013, which describe research in the life sciences; and (3) manuscripts from other journals matched for subject area and time of publication. We measured the change in the reporting of items included in the checklist.

## Identification of relevant manuscripts

### NPG publications

One individual was specifically employed by Nature to select studies which (a) described in vivo or in vitro research; (b) was published in Nature, Nature Neurology, Nature Immunology, Nature Cell Biology, Nature Chemical Biology, Nature Biotechnology, Nature Methods, Nature Medicine or Nature Structural and Molecular Biology. First, they identified papers accepted for publication with an initial submission date later than May 1st, 2013. Beginning with the then-current issues (volume corresponding to year 2015), they worked backwards in time, ensuring the submission date was after 1^st^ May 2013, collecting papers until they had 40 Nature papers and 20 each from other titles (“Post intervention” group). They then used a similar process to identify papers submitted for publication before 1st May 2013, matched for journal and for country of origin, starting with the May 2013 issue and working backwards, ensuring that the date of submission was after 1^st^ May 2011 (“pre-intervention” group). Where no match could be found with a submission date after 1st May 2011 (i.e. in a two year period) then the non-matched post intervention publication was excluded from analysis and a replacement post intervention publication selected, as above, with a matching pre-intervention publication then identified, as described above. Publications describing research involving only human subjects were not to be included. A Nature editorial administrator independent of publishing decisions reviewed manuscripts selection against the inclusion criteria and found some (less than10%) had been included incorrectly; they replaced these with manuscript pairs that they selected according to the inclusion algorithm. The published files corresponding to the publication pdfs (including the extended methods section, extended data and other supplementary materials) were used to generate pdfs for analysis. These were provided to a member of our research team (RM) at a different institution who used Adobe Acrobat to redact information relating to author names or affiliations, dates, volumes or page numbers; and the reference list; to minimise awareness of outcome assessors to whether the manuscript was pre‐ or post‐ intervention.

### Non‐ NPG publications

The same individual was responsible for identifying matching publications in other journals. They identified the NPG publication in PubMed and searched for “related citations” with the same calendar month of publication, selecting the first that was not published in an NPG Journal that also matched for whether it reported in vivo research, in vitro research, or both. If no matching related citation was found the extended the window of publication by 2 months, continuing until a matching publication was found. Because of a limited number of potential matching publications it was not possible to match non NPG manuscripts by country. The individual making this selection paid no further part in the study.

## Outcome assessment

The Nature checklist focussed on transparency in reporting and availability of materials and code, reflected in 10 items. We designed a series of questions (Appendix 1) to establish whether a given publication met or did not meet the requirements of the checklist. Where a manuscript described both in vivo and in vitro research, the series of questions was completed for each. Where there is more than one in vitro experiment or more than one in vivo experiment the question was considered in aggregate; that is, all experiments had to meet the requirements of the checklist item for it to be considered compliant.

Five researchers experienced in systematic review and risk of bias annotation scored the same 10 publications using our series of questions. Disagreements were resolved by group discussion, to arrive at a set of “Gold standard” answers for these 10 publications. We also used this experience to write a training guide for those seeking to use the checklist. We then used social media platforms and mailing lists to recruit outcome assessors. We had no prior requirements for the skills required of these individuals, but most had a background in medicine or biomedicine at graduate or undergraduate level; two were senior school students on Nuffield Research Placements in our group. After reviewing the training materials outcome assessors were invited sequentially to score publications from the “Gold standard” pool until their concordance with the Gold standard responses was 80% overall, and was 100% for the components of the primary outcome measure, for three successive publications. At this point we considered them to be trained. The training platform remains available for continuing professional development, at https://ecrf1.clinicaltrials.ed.ac.uk/npqip/Review/TrainingCover.

Pdf files of included manuscripts were uploaded to a bespoke website. Trained assessors were presented with manuscripts for scoring in random order. Each manuscript was scored by 2 individuals, one with experience in systematic review and risks of bias annotation and one recruited from outside this community. Disagreement between assessors were reconciled by a third, experienced individual who was not one of the original reviewers, who could see the responses previously given but not who the initial reviewers were.

### Statistical analysis plan

Given our focus on the reporting of measures to reduce the risks of bias we took as our primary outcome measure a composite measure of the proportion of publications meeting the relevant measures identified by Landis et al as being most important for transparency in reporting in vivo research. These are covered by items 2, 3 4 and 5 of the checklist and relate to the reporting of randomisation; of the blinded assessment of outcome; of sample size calculations; and of whether the manuscript described whether samples or animals were excluded from analysis. Importantly, checklist compliance did not require for example that the study was randomised; but rather that the authors stated whether or not it was randomised. The evaluation principle was to determine if someone with reasonable domain-knowledge could understand the parameters of experimental design sufficiently to inform interpretation. It has been argued that these measures might not be as relevant for exploratory studies, and for these we recorded the item as “not relevant”. We defined exploratory studies as those where hypothesis testing inferential statistical analyses were not reported. Where an item was not relevant for a publication (for instance with studies using transgenic animals where group allocation had been achieved by Mendelian randomisation) we considered compliance as meeting the remaining relevant criteria. Where a publication described both in vivo and in vitro experiments we analysed each type of experiment separately.

Our primary outcome was the change in the proportion of publications describing in vivo experiments published by NPG before and after May 2013 that meet all of the relevant Landis 4 criteria. We used the two-sample proportion test (prop.test) in R without the Yates continuity correction and two sided hypothesis testing to be sensitive to the possibility that performance might have declined rather than improved. Secondary outcomes were whether the proportion of publications describing in vivo experiments published by NPG after May 2013 which meet all four of the Landis 4 criteria was 80% or higher (Wald test; wald.ptheor.test, RVAideMemoire in R); the change in the proportion of publications describing in vitro experiments published by NPG before and after May 2013 which meet all four of the Landis 4 criteria (two sample proportion test as above); and the change of proportions in adequate reporting of statistical analysis details, individual Landis criteria, and descriptions of animals; reagents and their availability; sequence, structure or computer code deposition; and items relating to the involvement of human subjects or materials in included studies.. For the matching publications from non-NPG journals the secondary outcomes were the change in the proportion of publications describing in vivo experiments published before and after May 2013 which met all of the Landis 4 criteria (two sample proportion test); whether the proportion of publications describing in vivo experiments published after May 2013 which met all four of the Landis 4 criteria was 80% or higher (Wald test); the change in the proportion of publications describing in vitro experiments published before and after May 2013 which meet all four of the Landis 4 criteria 4 (two sample proportion test); and the change of proportions in adequate reporting of statistical analysis details, individual Landis criteria, and descriptions of animals; reagents and their availability; sequence, structure or computer code deposition; and items relating to the involvement of human subjects or materials in included studies. For each of these outcomes we compared the changes observed in NPG publications with that observed in non NPG publications. For each secondary analysis we used Holm Boneferroni correction using the p.adjust option for prop.test in R to account for the number of comparisons drawn, as described in Appendix B of the Data Analysis Plan. We also used interrupted time series analysis for each checklist item in an attempt to distinguish a discrete “shift” in performance from an upward “drift”, as described in the data analysis plan. A number of tertiary outcomes are described in the study protocol and statistical analysis plan and are reported in the supplementary material.

### Power Calculations

In planning the study we performed power calculations in STATA. The power to detect changes in reporting depended on the baseline performance; with baseline prevalence of compliance of 10% we had 80% power to detect an absolute increase of 13% to 23% at a significance level of p<0.01; with baseline compliance of 50% we had 80% power to detect an absolute increase of 16% to 66% at a significance level of p<0.01. For secondary outcomes we had lower statistical power, but after correction for the number of comparisons made we had at worse 67% power to detect a 15% improvement in the reporting of any individual item.

## Results

896 publications were identified and uploaded for outcome ascertainment, 448 in each cohort. 2 non-NPG manuscripts were excluded because they did not meet the inclusion criteria, and we identified 4 NPG and 9 non-NPG manuscripts included more than once. 444 NPG publications and 437 non-NPG publications underwent outcome assessment. One NPG publication and one non-NPG publication were adjudged at the time of outcome assessment to report neither in vivo nor in vitro research and so were excluded. The analysis is therefore based on 443 NPG publications (219 before and 224 after 1^st^ May 2013) and 436 non-NPG publications (194 before and 242 after 1^st^ May 2013) (Figure1). The difference in numbers for NPG and non-NPG before and after 1^st^ May 2013 is because some of the NPG “before” papers matched best with publications in other journals published in the few months following May 2013. Overall, 43% of matched pairs had dates of publication within 1 month, 54% within 2 months, 64% within 3 months and 81% within 6 months of each other (range −11 to +22 months). 239 publications described only in vivo research, 132 described only in vitro research, and 508 described both. The source journals are given in Table 1; in total 198 different titles contributed matching publications (median manuscripts per publication 1, range 1 - 47). The PMIDs of included publications are listed in the data supplement.

**Figure 1:**
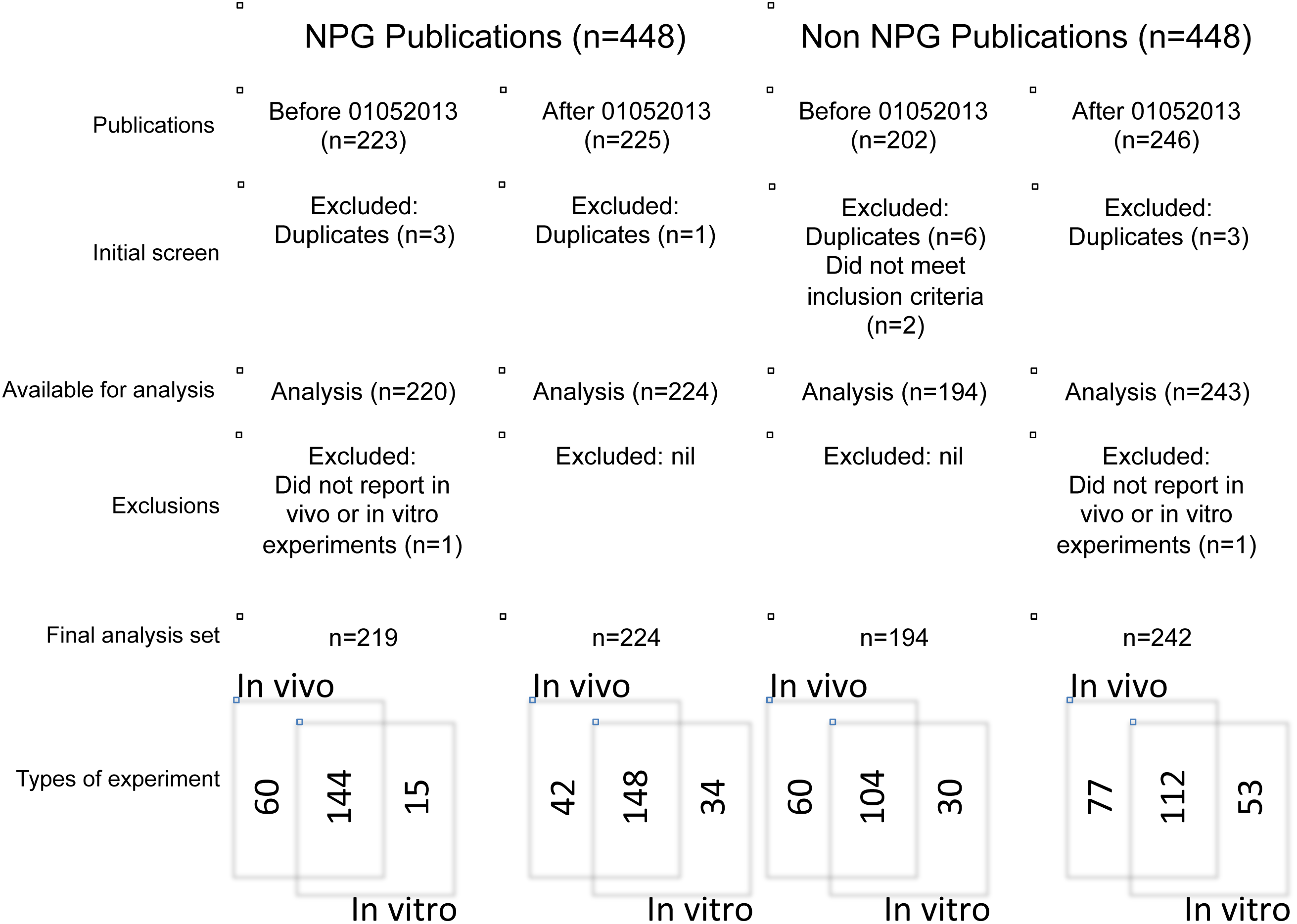
Manuscripts initially included, and reasons for exclusion, and type of experiments described.

**Table 1:** Sources of manuscripts included in the study

| **Journal** | **number** |
| --- | --- |
| Nature | 89 |
| Nat Neurosci | 45 |
| Nat Med | 44 |
| Nat Immunol | 44 |
| Nat Cell Biol | 44 |
| Nat Methods | 43 |
| Nat Genet | 40 |
| Nat Biotechnol | 40 |
| Nat Chem Biol | 35 |
| Nat Struct Mol Biol | 19 |
| PLoS One | 47 |
| Proc Natl Acad Sci U S A | 24 |
| J Neurosci | 19 |
| J Biol Chem | 13 |
| J Immunol | 13 |
| Other | 320 |

205 individuals registered with the project, of whom 38 completed their training and 35 assessed at least one manuscript. 12 also served as reconcilers, and the web interface was programmed to ensure that they were not offered for reconciliation manuscripts that they had previously adjudicated. Including reconciliation, the median number of manuscripts scored was 13 (range 1 to 441). The agreement between the initial pair of outcome assessors ranged from being no better than chance at 50% (in vivo studies, Implementation of statistical methods and measures: “Is the variance similar (difference less than two-fold) between the groups that are being statistically compared?”) to 98% (in vivo studies, “Does the study report the species?”). Median agreement was 82%. (IQR 68 - 89%).

### Reporting of the Landis 4 items

The proportion of NPG in vivo studies reaching full compliance with the Landis 4 criteria increased from 0% (0/204) to 16.3% (31/190) (X^2^ = 36.1, df = 1, p = 1.8x10-^9^), but remained significantly lower than the target of 80% (95% CI 11.7% to 22.3%, Wald test v 80% t = -15.4, p = 2.2x10-^16^).

For randomisation to experimental group, the preferred standard is that the manuscript describes which method of randomization was used to determine how samples or animals were allocated to experimental groups, although manuscripts were also compliant if they included a statement about randomization even if no randomization was used. The proportion of NPG in vivo studies reporting randomisation was 1.8% (3/170, 95% CI 0.6 to 5.3%) before and 11.2% (19/170, 95% CI 7.2 to 16.9%) after (**x^2^** = 12.4, df = 1, adj p = 0.054). The proportion of studies mentioning randomisation even where it was not reported increased from 8.3% (14/169, 95% CI 5.0 to 13.5) to 64.2% (97/151, 95% CI 56.3 to 71.5%)(x^2^ = 110.2, df = 1, adj p = 3.2x10-^14^). Figure 2(a) shows change in the proportion of studies meeting these criteria before and after the change in editorial policy.

**Figure 2:**
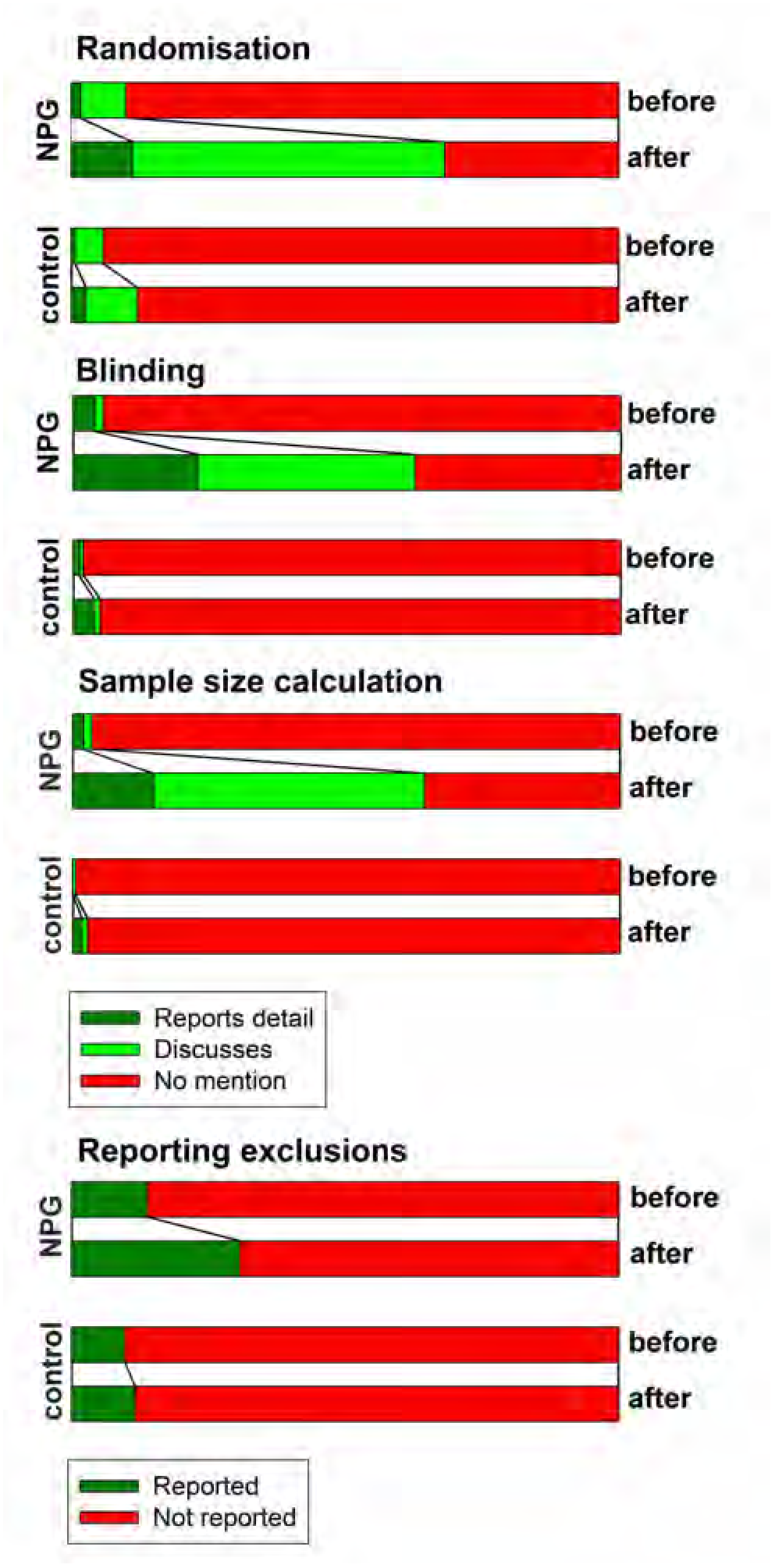
Compliance with each Landis criteria for in vivo experiments for NPG and non NPG manuscripts before and after 1^st^ May 2013.

For blinding, the preferred standard is that the manuscript describes whether the investigator was blinded to the group allocation during the experiment and/or when assessing the outcome, although manuscripts were also compliant if they included a statement about blinding even if no blinding was done. The proportion of NPG in vivo studies reporting blinding during group allocation or outcome assessment or both increased from 4% (8/198, 95% CI 2.0 to 7.9%) to 22.8% (42/184, 95% CI 17.3 to 29.4%)(X^2^ = 29.6, df = 1, adj p = 7.6x10-^6^). The proportion of studies mentioning blinding even where it was not reported increased from 1.6% (3/182, 95% CI 0.5 to 5.0%) to 55.3% (73/132, 95% CI 46.8 to 65.6%)(X^2^ = 120.1, df = 1, adj p < 3.2x10-^14^). Figure 1(b) shows change in the proportion of studies meeting these criteria before and after the change in editorial policy.

The proportion of studies reporting animals excluded from analysis increased from 13.9% (28/202, 95% CI 9.7 to 19.3%) to 30.7% (58/189, 95% CI 24.5 to 36.7%)(X^2^; = 16.1, df = 1, adj p = 0.008). Figure 1(c) shows change in the proportion of studies meeting these criteria before and after the change in editorial policy.

For sample size calculations, the preferred standard is that the manuscript describes how the sample size was chosen to ensure adequate power to detect a pre-specified effect size, although manuscripts were also compliant if they included a statement about sample size estimate even if no statistical methods were used. The proportion of studies reporting an a priori sample size calculation increased from 2.0% (4/196, 95% CI 0.8 to 5.3%) to 14.8% (27/182, 95% CI 10.4 to 20.8%)(X^2^ = 20.5, df = 1, adj p = 0.0008). The proportion of studies mentioning sample size even where a sample size calculation was not reported increased from 1.6% (3/192, 95% CI 0.5 to 4.7%) to 58.4% (90/154, 95% CI 50.5 to 66.0%)(X^2^ = 140.7, df = 1, adj p < 3.2x10^-14^). Figure 1(d) shows change in the proportion of studies meeting these criteria before and after the change in editorial policy.

For NPG in vitro studies, the proportion reaching full compliance with the Landis 4 criteria was 0% (0/159) before and 3.3% (6/176) after (X^2^ = 6.8, df = 1, Holm Bonferroni adjusted p = 1.00). The proportion of studies reporting randomisation was 0% (0/149) before and 2.9% (5/173, 95% CI 1.2 to 6.8%) after (X^2^ = 4.4, df = 1, adj p=1.00). The proportion of studies mentioning randomisation even where it was not reported increased from 0% (0/149) to 15.6% (97/151, 95% CI 10.8 to 21.9%)(X^2^ = 25.3, df = 1, p = 6.9x10^-5^). The proportion of studies reporting blinding during group allocation or outcome assessment or both was 3.9% (6/155, 95% CI 1.8 to 8.4%) before and 8.9% (16/179, 95% CI 5.6 to 14.1) after (X^2^; = 3.467, df = 1, p=1.00). The proportion of studies mentioning blinding even where it was not reported increased from 0.7% (1/150, 95% CI 0.1 to 4.6%) to 15.9% (25/157, 95% CI 11.0 to 22.5) (X^2^; = 23.0, df = 1, p=0.0002). The proportion of studies reporting exclusions from analysis was 8.2% before (13/159, 95% CI 4.8 to 13.6%) and 15.9% (29/182, 95% CI 11.3 to 22.0%) after (X^2^ = 4.73, df = 1, p =1.00). The proportion of studies reporting an a priori sample size calculation was 1.3% (2/155, 95% CI 0.3 to 5.0%) before and 7.9% (14/177, 95% CI 5.1 to 13.5%) after (X^2^ = 8.7106, df = 1, p = 1.00). The proportion of studies mentioning sample size even where a sample size calculation was not reported increased from 3.3% (5/153, 95% CI 1.4 to 7.6%) to 28.5% (47/165, 95% CI 22.1 to 35.8%)(X^2^=36.9, df = 1, p=1.8x10^-7^).

The proportion of matching (non-NPG) in vivo studies reaching full compliance with the Landis 4 criteria was 1% (1/164) before and 1% (1/189) after (X^2^ = 0.01, df = 1, adj p = 1.00), and for in vitro studies, the proportion of non-NPG studies reaching full compliance with the Landis 4 criteria was 0% (0/134) before and 1% (1/165) after (X^2^ = 0.8, df = 1, adj p = 1.00). The prevalence of reporting the different items before and after is shown in table 2; there was no significant change in reporting of any of the individual Landis 4 criteria for either in vivo or in vitro research.

**Table 2:** Primary outcome: Compliance with Landis 4 guidelines, in vivo research.

| Item | NPG before |  |  |  | NPG after |  |  |  | p | Matched before |  |  |  | Matched after |  |  |  | p |
| --- | --- | --- | --- | --- | --- | --- | --- | --- | --- | --- | --- | --- | --- | --- | --- | --- | --- | --- |
|  | n | N | % | CI | n | N | % | CI |  | n | N | % | CI | n | N | % | CI |  |
| In vivo |  |  |  |  |  |  |  |  |  |  |  |  |  |  |  |  |  |  |
| Full Landis | 0 | 204 | 0 | 0.0-2.3 | 31 | 190 | 16.3 | 11.7-22.3 | <10<sup>-8</sup> | 1 | 164 | 0.6 | 0.1-4.2 | 1 | 189 | 0.5 | 0.1-3.7 | n.s. |

Legend: n, number meeting criteria: N, total number of studies: %, percent meeting criteria: CI, 95% confidence interval of that percentage: p, significance level (two sample proportion test): n.s., not significant at p<0.05.

**Table 3:** Secondary outcome: Full Landis compliance, in vitro research.

| Item | NPG before |  |  |  | NPG after |  |  |  |  | Matched before |  |  |  | Matched after |  |  |  |  |
| --- | --- | --- | --- | --- | --- | --- | --- | --- | --- | --- | --- | --- | --- | --- | --- | --- | --- | --- |
|  | n | N | % | CI | n | N | % | CI | Adj p | n | N | % | CI | n | N | % | CI | Adj p |
| In vitro |  |  |  |  |  |  |  |  |  |  |  |  |  |  |  |  |  |  |
| Full Landis | 0 | 159 | 0.0 | 0.0-2.3 | 6 | 182 | 3.3 | 1.5-7.1 | n.s. | 0 | 134 | 0.0 | 0.0-3.5 | 1 | 165 | 0.6 | 0.1-4.2 | n.s. |

Legend: n, number meeting criteria: N, total number of studies: %, percent meeting criteria: CI, 95% confidence interval of that percentage: Adj p, adjusted significance level (two sample proportion test (prop.test) followed by Holm Bonferroni correction (p.adjust.methods)): n.s., not significant at $p < 0.05$ .

**Table 4:** Compliance with individual Landis 4 items, in vivo and in vitro research.

| Item | NPG before |  |  |  | NPG after |  |  |  |  | Matched before |  |  |  | Matched after |  |  |  |  |
| --- | --- | --- | --- | --- | --- | --- | --- | --- | --- | --- | --- | --- | --- | --- | --- | --- | --- | --- |
|  | n | N | % | CI | n | N | % | CI | Adj p | n | N | % | CI | n | N | % | CI | Adj p |
| In vivo |  |  |  |  |  |  |  |  |  |  |  |  |  |  |  |  |  |  |
| Report method of rand? | 3 | 170 | 1.8 | 0.6-5.3 | 19 | 170 | 11.2 | 7.2-16.9 | n.s. | 1 | 134 | 0.8 | 0.1-5.1 | 4 | 149 | 2.7 | 1.0-6.9 | n.s. |
| statement about randomisation | 14 | 169 | 8.3 | 5.0-13.5 | 97 | 151 | 64.2 | 56.3-71.5 | 3x10<sup>-14</sup> | 7 | 136 | 5.1 | 2.5-10.4 | 14 | 147 | 9.5 | 5.6-15.4 | n.s. |
| Blinded? | 8 | 198 | 4.0 | 2.0-7.9 | 42 | 184 | 22.8 | 17.3-29.4 | 8x10<sup>-6</sup> | 2 | 162 | 1.2 | 0.3-4.8 | 7 | 183 | 3.8 | 1.8-7.8 | n.s. |
| Statement about blinding | 3 | 182 | 1.6 | 0.5-5.0 | 73 | 132 | 55.3 | 46.8-63.4 | 3x10<sup>-14</sup> | 1 | 151 | 0.7 | 0-4.6 | 2 | 165 | 1.2 | 0-4.7 | n.s. |
| Exclusions reported? | 28 | 202 | 13.9 | 9.7-19.3 | 58 | 189 | 30.7 | 24.5-37.6 | 0.008 | 16 | 164 | 9.8 | 6.1-15.3 | 22 | 189 | 11.6 | 7.8-17.0 | n.s. |
| Exclusion criteria defined? | 24 | 200 | 12 | 8.2-17.3 | 35 | 188 | 18.6 | 13.7-24.8 | n.s. | 14 | 163 | 8.6 | 5.2-14.0 | 20 | 188 | 10.6 | 7.0-15.9 | n.s. |
| Clear these were pre-specified? | 1 | 25 | 4 | 0.6-23.6 | 5 | 39 | 12.8 | 5.4-27.3 | n.s. | 0 | 17 | 0 | 0.0-22.9 | 0 | 21 | 0 | 0.0-19.2 | n.s. |
| Was a SSC done? | 4 | 196 | 2.0 | 0.8-5.3 | 27 | 182 | 14.8 | 10.4-20.8 | 0.0008 | 0 | 156 | 0 | 0.0-3.0 | 3 | 183 | 1.6 | 0.5-5.0 | n.s. |
| If not done, was SSC mentioned? | 3 | 192 | 1.6 | 0.5-4.7 | 90 | 154 | 58.4 | 50.5-66.0 | 3x10<sup>-14</sup> | 1 | 157 | 0.6 | 0-4.4 | 2 | 180 | 1.1 | 0.3-4.3 | n.s. |
| In vitro |  |  |  |  |  |  |  |  |  |  |  |  |  |  |  |  |  |  |
| Report method of rand? | 0 | 149 | 0 | 0.0-3.1 | 5 | 173 | 2.9 | 1.2-6.8 | n.s. | 1 | 125 | 0.8 | 0.1-5.5 | 1 | 157 | 0.6 | 0.1-4.4 | n.s. |
| statement about randomisation | 0 | 149 | 0 | 0.0-3.1 | 26 | 167 | 15.6 | 10.8-21.9 | 7x10<sup>-5</sup> | 0 | 123 | 0 | 0.0-3.8 | 4 | 156 | 2.6 | 1.0-6.6 | n.s. |
| Blinded? | 6 | 155 | 3.9 | 1.8-8.4 | 16 | 179 | 8.9 | 5.6-14.1 | n.s. | 3 | 131 | 2.3 | 0.7-6.9 | 1 | 162 | 0.6 | 0.1-4.2 | n.s. |
| Statement about blinding | 1 | 150 | 0.7 | 0.1-4.6 | 25 | 157 | 15.9 | 11.0-22.5 | 0.0002 | 0 | 127 | 0 | 0.0-3.7 | 1 | 158 | 0.6 | 0.1-4.3 | n.s. |
| Exclusions reported? | 13 | 159 | 8.2 | 4.8-13.6 | 29 | 182 | 15.9 | 11.3-22.0 | n.s. | 7 | 133 | 5.3 | 2.5-10.6 | 10 | 165 | 6.1 | 3.3-10.9 | n.s. |
| Exclusion criteria defined? | 12 | 159 | 7.5 | 4.3-12.8 | 23 | 178 | 12.9 | 8.7-18.7 | n.s. | 7 | 133 | 5.3 | 2.5-10.6 | 9 | 165 | 5.5 | 2.9-10.2 | n.s. |
| Clear these were pre-specified? | 0 | 14 | 0 | 0.0-26.8 | 1 | 24 | 4.2 | 0.6-24.4 | n.s. | 0 | 8 | 0 | 0.0-40.0 | 0 | 11 | 0 | 0.0-32.1 | n.t. |
| Was a SSC done? | 2 | 155 | 1.3 | 0.3-5.0 | 14 | 177 | 7.9 | 5.1-13.5 | n.s. | 0 | 129 | 0 | 0.0-3.6 | 0 | 161 | 0 | 0.0-2.9 | n.s. |
| If not done, was SSC mentioned? | 5 | 153 | 3.3 | 1.4-7.6 | 47 | 165 | 28.5 | 22.1-35.8 | 2x10<sup>-7</sup> | 0 | 129 | 0 | 0.0-3.6 | 1 | 162 | 0.6 | 0.1-4.2 | n.s. |

Legend: n, number meeting criteria: N, total number of studies: %, percent meeting criteria: CI, 95% confidence interval of that percentage: Adj p, adjusted significance level (two sample proportion test (prop.test) followed by Holm Bonferroni correction (p.adjust.methods)): n.s., not significant at p<0.05: n.t. not tested (n<10 for one of the comparisons)

**Table 5:**
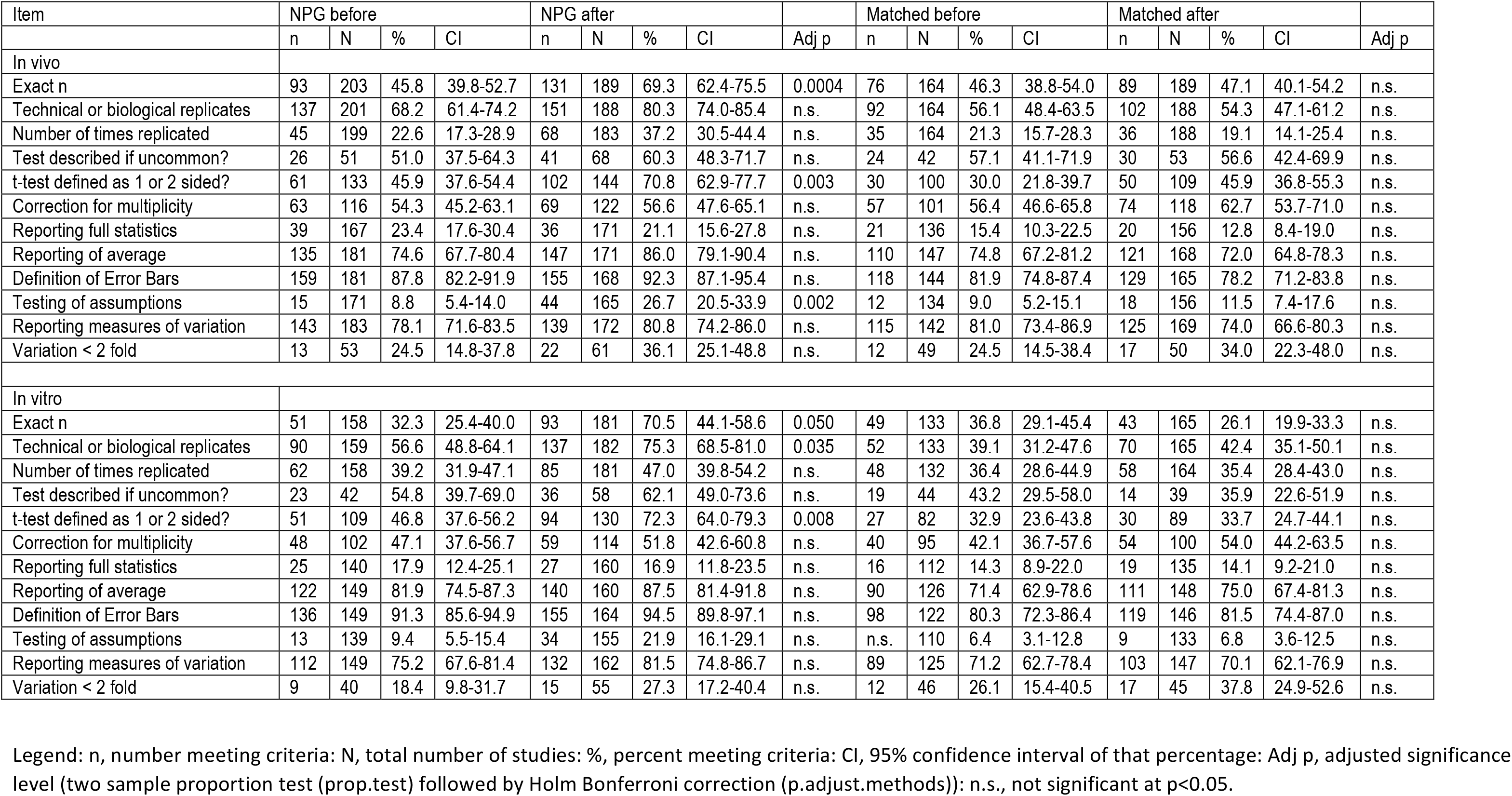
Secondary outcome: statistical items.

**Table 6:** Other secondary outcomes.

| Item | NPG before |  |  |  | NPG after |  |  |  |  | Matched before |  |  |  | Matched after |  |  |  |  |
| --- | --- | --- | --- | --- | --- | --- | --- | --- | --- | --- | --- | --- | --- | --- | --- | --- | --- | --- |
|  | n | N | % | CI | n | N | % | CI | Adj p | n | N | % | CI | n | N | % | CI | Adj p |
| Animals |  |  |  |  |  |  |  |  |  |  |  |  |  |  |  |  |  |  |
| Was the species reported? | 203 | 203 | 100 | 97.7-100 | 189 | 189 | 100.0 | 97.5-100 | n.s. | 163 | 164 | 99.4 | 95.8-99.9 | 188 | 189 | 99.5 | 96.3-99.9 | n.s. |
| Was the strain reported? | 187 | 203 | 92.1 | 87.5-95.1 | 181 | 189 | 95.8 | 91.8-97.9 | n.s. | 149 | 164 | 90.9 | 85.4-94.4 | 176 | 189 | 93.1 | 88.5-96.0 | n.s. |
| Was the sex reported? | 69 | 193 | 35.8 | 29.3-47.8 | 96 | 184 | 52.2 | 45.0-59.3 | n.s. | 59 | 161 | 36.6 | 29.6-44.4 | 67 | 183 | 36.3 | 30.0-43.8 | n.s. |
| Was exact age or weight given? | 31 | 203 | 15.3 | 11.0-20.9 | 41 | 188 | 21.8 | 16.5-28.3 | n.s. | 41 | 164 | 25.0 | 19.0-32.2 | 39 | 189 | 20.6 | 15.4-27.0 | n.s. |
| Was ethical approval reported? | 116 | 194 | 59.8 | 52.7-66.5 | 121 | 178 | 68.0 | 60.1-74.4 | n.s. | 91 | 151 | 60.3 | 52.3-67.8 | 123 | 180 | 68.3 | 61.2-74.7 | n.s. |
| Ethical guidelines reported? | 148 | 199 | 74.4 | 67.9-80.0 | 154 | 181 | 85.1 | 79.1-89.6 | n.s. | 111 | 151 | 73.5 | 65.9-79.9 | 137 | 181 | 75.7 | 68.9-81.4 | n.s. |
| Reagents |  |  |  |  |  |  |  |  |  |  |  |  |  |  |  |  |  |  |
| In vivo Antibodies | 89 | 142 | 62.7 | 54.4-70.2 | 98 | 135 | 72.6 | 64.5-79.4 | n.s. | 38 | 115 | 33.0 | 25.1-42.1 | 58 | 127 | 45.7 | 37.2-54.4 | n.s. |
| In vitro antibodies | 75 | 125 | 60.0 | 51.2-68.2 | 107 | 142 | 75.4 | 67.6-81.7 | n.s. | 29 | 104 | 27.9 | 20.1-37.2 | 51 | 126 | 40.5 | 32.2-49.2 | n.s. |
| Total antibodies | 164 | 267 | 61.4 | 55.4-67.1 | 205 | 277 | 74.0 | 68.5-78.8 | n.s. | 67 | 219 | 30.6 | 24.8-37.0 | 109 | 253 | 43.1 | 37.1-49.3 | n.s. |
| In vitro: cell line source | 51 | 102 | 50.0 | 40.4-59.6 | 96 | 137 | 70.1 | 61.9-77.1 | n.s. | 53 | 84 | 63.1 | 52.3-72.7 | 62 | 111 | 55.9 | 46.5-64.8 | n.s. |
| Recent authentication? | 1 | 95 | 1.1 | 0.2-7.1 | 9 | 126 | 7.1 | 3.8-13.2 | n.s. | 4 | 76 | 5.3 | 2.0-13.2 | 1 | 97 | 1.0 | 0.2-7.0 | n.s. |
| Recent mycoplasma testing? | 1 | 97 | 1.0 | 0.2-7.0 | 33 | 127 | 26.0 | 19.1-34.3 | 4x10<sup>-5</sup> | 1 | 77 | 1.3 | 0.2-8.6 | 1 | 97 | 1.0 | 0.2-7.0 | n.s. |
| Accession: DNA/protein | 30 | 61 | 49.2 | 36.9-61.5 | 32 | 64 | 50.0 | 38.0-62.0 | n.s. | 10 | 21 | 47.6 | 27.8-68.2 | 19 | 45 | 42.2 | 28.8-56.9 | n.s. |
| Accession: Macromolecular | 0 | 4 | 0 | 0.0-60.4 | 4 | 7 | 57.1 | 20.2-88.2 | n.t. | 0 | 2 | 0 | 0.0-80.2 | 3 | 4 | 75.0 | 21.9-98.7 | n.t. |
| Accession: Crystallography | 5 | 7 | 71.4 | 30.2-94.9 | 3 | 11 | 27.3 | 7.3-60.7 | n.t. | 1 | 2 | 50.0 | 9.4-90.5 | 0 | 1 | 0 | 0.0-94.5 | n.t. |
| Accession: Microarray | 12 | 33 | 36.4 | 21.9-53.7 | 21 | 38 | 55.3 | 39.5-70.1 | n.s. | 7 | 15 | 46.7 | 24.1-70.7 | 12 | 18 | 66.7 | 42.9-84.2 | n.s. |
| Accession: Other | 2 | 7 | 28.66 | 7.2-67.3 | 8 | 18 | 44.4 | 24.0-67.0 | n.t. | 1 | 5 | 20.0 | 10.5-70.1 | 4 | 6 | 66.7 | 24.1-94.0 | n.t. |
| Computer Code with paper? | 3 | 14 | 21.4 | 7.1-49.4 | 5 | 24 | 20.8 | 9.0-41.3 | n.s. | 0 | 5 | 0 | 0.0-53.7 | 2 | 14 | 14.3 | 3.6-42.7 | n.t. |
| Code in public domain? | 3 | 11 | 27.3 | 9.0-58.6 | 5 | 24 | 20.8 | 9.0-41.3 | n.s. | 0 | 5 | 0 | 0.0-53.7 | 2 | 13 | 15.4 | 3.9-45.1 | n.t. |
| Was that code accessible? | 2 | 3 | 66.7 | 12.5-98.2 | 5 | 7 | 71.4 | 30.2-94.9 | n.t. | 0 | 1 | 0 | 0.0-94.5 | 2 | 4 | 50.0 | 15.0-85.0 | n.t. |
| Did the code function ? | 2 | 3 | 67.7 | 12.5-98.2 | 2 | 2 | 100.0 | 19.8-100 | n.t. |  | 0 |  |  |  | 0 |  |  | n.t. |
| Say where you could get code? | 0 | 9 | 0 | 0.0-37.1 | 1 | 16 | 6.3 | 0.3-32.3 | n.t. | 2 | 5 | 40.0 | 7.2-83.0 | 0 | 10 | 0 | 0.0-34.4 | n.t. |
| Human materials |  |  |  |  |  |  |  |  |  |  |  |  |  |  |  |  |  |  |
| Reporting ethical approvals | 35 | 43 | 81.4 | 67.0-90.4 | 46 | 56 | 82.1 | 69.9-90.1 | n.s. | 13 | 23 | 56.5 | 36.3-74.8 | 29 | 38 | 76.3 | 60.4-87.2 | n.s. |
| Reporting consent | 35 | 42 | 83.3 | 69.0-91.8 | 47 | 56 | 83.9 | 71.9-91.4 | n.s. | 15 | 24 | 62.5 | 42.2-79.2 | 24 | 38 | 63.2 | 47.0-76.8 | n.s. |
| Consent to photos | 2 | 3 | 66.7 | 12.5-98.2 | 1 | 1 | 100.0 | 5.4-100 | n.t. |  | 0 |  |  | 0 | 2 | 0 | 0.0-80.2 | n.t. |
| Clinical trial number |  | 0 |  |  | 2 | 3 | 66.7 | 12.5-98.2 | n.t. | 1 | 1 | 100.0 | 5.4-100 |  | 0 |  |  | n.t. |
| CONSORT |  | 0 |  |  |  | 0 |  |  | n.t. | 0 | 1 | 0 | 0.0-94.5 |  | 0 |  |  | n.t. |
| REMARK | 0 | 1 | 0 | 0.0-94.5 |  | 0 |  |  | n.t. | 0 | 2 | 0 | 0.0-80.2 | 0 | 1 | 0 | 0.0-94.5 | n.t. |

Legend: n, number meeting criteria: N, total number of studies: %, percent meeting criteria: CI, 95% confidence interval of that percentage: Adj p, adjusted significance level (two sample proportion test (prop.test) followed by Holm Bonferroni correction (p.adjust.methods)): n.s., not significant at p<0.05: n.t. not tested (n<10 for one of the comparisons).

### Statistical reporting

For in vivo studies reported in NPG manuscripts there were significant improvements in the reporting of exact numbers (from 46% to 69%), of whether t-tests were defined as one or two sided (from 46% to 71%), and whether the assumptions of the test had been checked (from 9% to 27%). For in vitro experiments described in NPG manuscripts there were significant improvements in the reporting of the exact numbers (from 32% to 70%); of whether data represented technical or biological replicates (from 57% to 75%); and whether t-tests were defined as one or two sided (from 47% to 72%). For in vivo and in vitro studies described in non-NPG publications there was no significant change in any of the items relating to statistical reporting.

### Other checklist items

For reporting of details of animals used, reporting of animal species and strain was high even before the change in editorial policy. There was no significant change in reporting any of these items in NPG‐ and non-NPG manuscripts, or in the reporting of details of antibodies used. For in vitro research, there was an increase in the proportion of studies in NPG manuscripts reporting recent mycoplasma testing of the cell lines used (from 1% to 26%) but not for non-NPG manuscripts (1% before, 1% after). For reporting and availability of accession data (eg DNA or protein sequence deposition) and computer code there were no significant changes for either NPG or non-NPG publications. Finally, there were no significant changes in the reporting of items relating to human subjects or the use of human materials, but for most items the number of publications for which these were relevant was very low indeed.

We were also interested in whether changes in reporting had occurred as a step change at the time of the change in editorial policy; whether there was an initial improvement with then a return to previous performance; or if there was an ongoing improvement in reporting. To address these we conducted an interrupted time series analysis, to estimate the rate of change before the intervention; any step change at the time of the intervention; and the rate of change after the intervention. We grouped publications in 3 month periods starting November 2011, and for each quarter calculated the proportional compliance with the criteria in question. Because publications were not evenly distributed across time the analysis is of substantially reduced power, but the fitted lines for overall compliance and for each component of the Landis checklist for in vivo research are shown in Figure 3. It appears that with the exception of sample size calculation there is a continuing improvement over time in both NPG and non NPG publications; for sample size calculations the improvement is only seen in NPG publications. Figure 4 shows radar charts of compliance for each checklist item in NPG ans non NPG manuscripts before and after May 2013.

**Figure 3:**
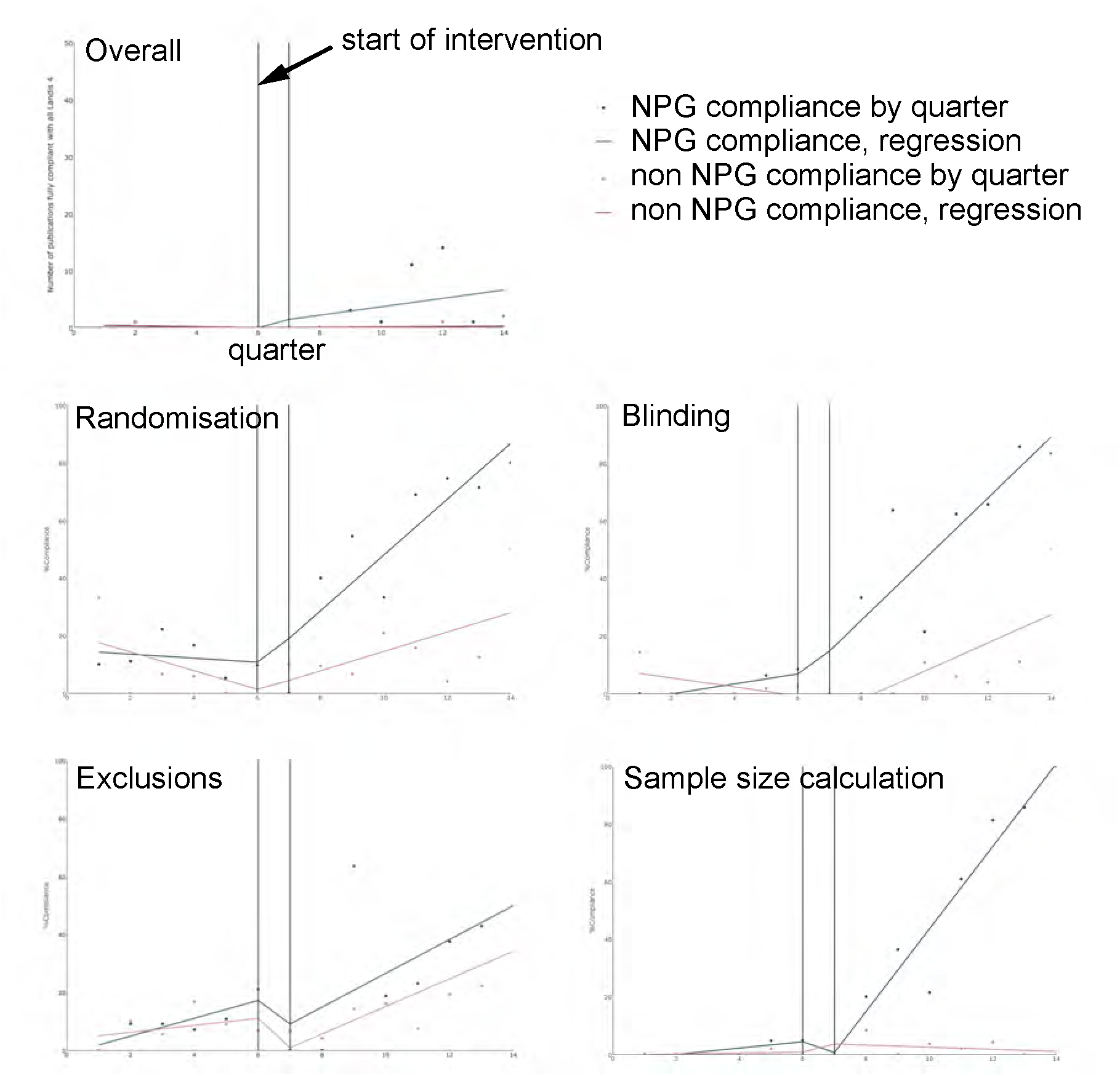
Interrupted time series analysis for overall Landis compliance and compliance with Landis components in in vivo experiments reported in NPG and non NPG manuscripts. Quarter 6 began on 1^st^ May 2013.

**Figure 4:**
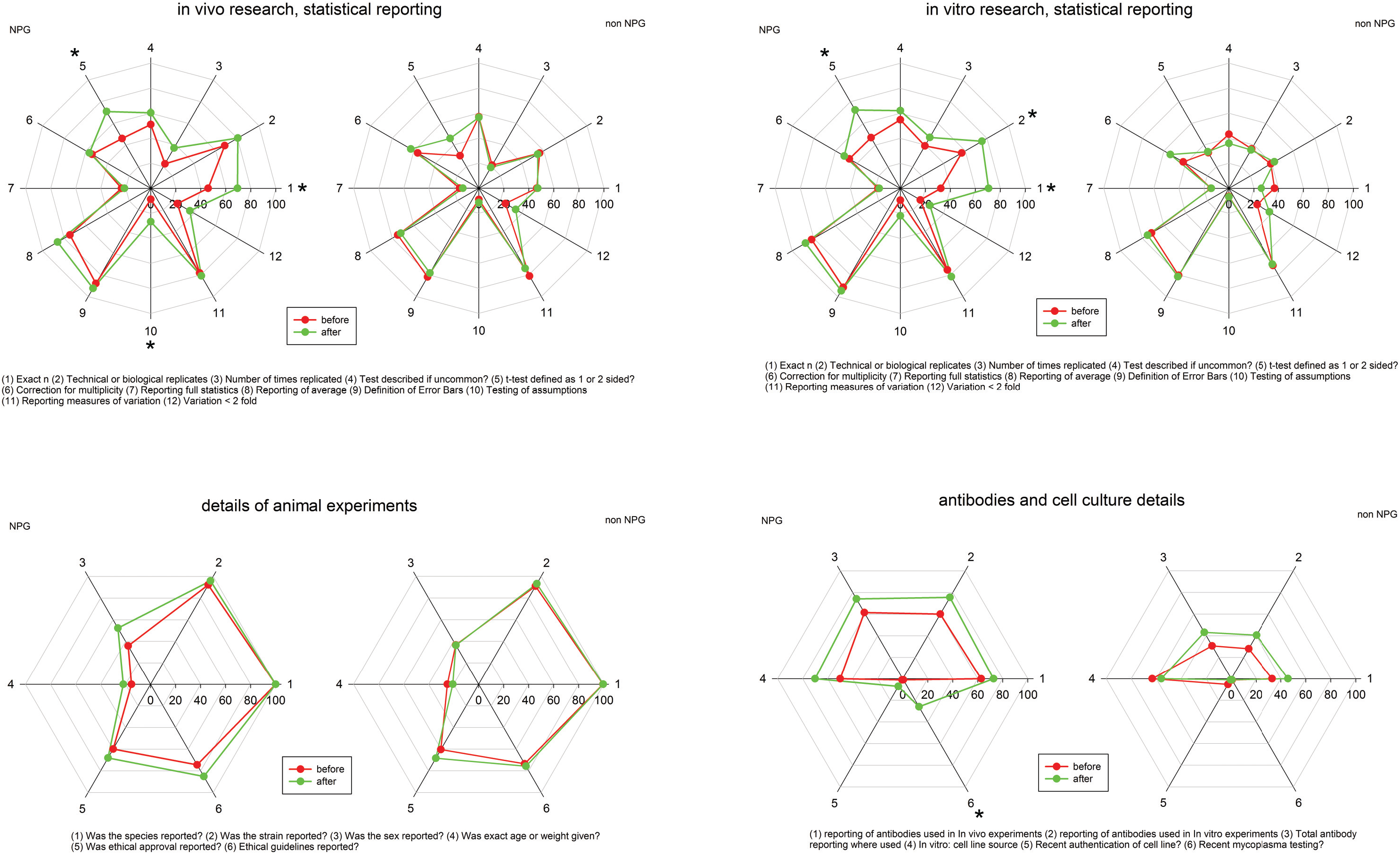
Radar plots for compliance with individual components of the NPG checklist before (red) and after (green) 1^st^ May 2013 for (a) statistical reporting, in vivo research; (b) statistical reporting, in vitro research; (c) reporting of details of animals used; and (d) reporting of reagents used. * adjusted p < 0.05 for change between “before” and “after”.

## Discussion

The change in editorial policy at NPG was associated with major improvements in reporting of randomisation, blinding, exclusions from analysis and sample size calculations. For the highly challenging primary outcome measure, full compliance increased from zero to 16%. This falls short of the target compliance of 80%, but should be seen in the context firstly that only 1 of 1073 publications from 2009-10 from leading UK institutions achieved this standard[8]; and secondly that overall compliance of 80% would require compliance with individual items of around 95%.

The checklist relates to transparency in reporting, and manuscripts were judged to be compliant if they either reported measures to address that risk of bias, or reported that such measures were not taken. For reports of in vivo research, compliance for randomisation, blinding, reporting of exclusions and sample size calculations in NPG publications reached 68%, 62%, 31% and 64% respectively. For non NPG publications the performance was 12%, 5%, 12% and 3%. The figures for NPG publications are similar to those recently reported for in vivo research published in the journal “Stroke” [9], which began requiring reporting of such details following the publication of good practice guidelines in 2009 [10]; and where performance was found to be substantially higher than for in vivo research published in other American Heart Association journals.

For reports of in vitro research, compliance was substantially lower. There have been few systematic attempts to measure the quality of reporting of measures to reduce the risks of bias in vitro research, and our findings suggest that, both in NPG and non NPG journals, this remains low. There were improvements in reporting randomisation, blinding and sample size calculations in NPG descriptions of in vitro research, but only to 18%, 23% and 34% respectively. For non NPG the equivalent figures were 3%, 1% and 1%. There were no significant changes in the reporting of exclusion of in vitro data, with post intervention compliance of 16% in NPG publications and 6% in non NPG publications.

For other checklist items, changes in performance were less dramatic, but there appeared to be incremental improvements across most of the items measured, although few of these breached our rather parsimonious adjustment for multiple testing. In spite of substantial attention given to the importance of reporting the sex of experimental animals this was only done in 52% of post intervention NPG studies and in 36% of non NPG studies.

Ours is an observational study, and it is possible that other (related or unrelated) changes were responsible for much if not all of the differences seen. These changes were not observed at other journals (at least not when taken in aggregate), and so it is likely that alternative causal factors would relate to NPG editorial policy and practice. While we are not aware of any other relevant changes in editorial policy occurring at a relevant time, it is likely that this change in editorial policy was accompanied by increased attention given to the importance of the quality of reporting by both in house editorial staff and external peer reviewers. It is not possible to determine whether these might have caused the changes seen. However, a randomised controlled study of the effect of ARRIVE checklist completion on the quality of reporting of in vivo research at PLoS One will report shortly.

During the course of the study we encountered some difficulties that we had not expected. We had thought that it would be straightforward to distinguish between an in vivo experiment and an in vitro experiment, but we had to develop an operational approach which defined that experiment on the basis of the subject at the time that the experimental intervention occurred; so a tissue slice experiment involving tissues from animals exposed to treatment or control we considered in vivo; while a similar experiment applying drugs directly to the slice we considered to be an in vitro experiment.

Further, there were some checklist items where agreement between outcome assessors was very low – for instance, for the question of whether for in vivo research the difference in variance between groups being compared was less than two fold, the agreement was no better than would be expected by chance alone. We recommend that the development of publication checklists should include an assessment of inter-observer variation by potential users of the checklist for each checklist item; low agreement might indicate that the item should be rephrased or reframed, or that more explanatory text is required.

Finally, our work shows the challenge of assessing even a relatively limited number of publications against a relatively straightforward checklist. We are delighted that so many collaborators (from 6 continents) agreed to participate, and are very grateful to them. However, even with their help the outcome assessment and reconciliation took 17 months. This is too slow to be useful for instance for quality improvement activity, where more rapid feedback would allow more rapid adjustments in response to performance. We have tested the use of text analytics using regular expressions to automatically ascertain reporting of measures to reduce the risk of bias, and for some such risks of bias the approach achieves sensitivities and specificities above 80%. However, for more complex items it may be that machine learning approaches using for instance convoluted neural networks may be more successful, and this is a current focus of our research. We hope that, by making the dataset for this study available, this might be used for instance for distant supervised learning in such systems.

## Conclusions

Introduction of a checklist lead to substantial improvements in the quality of reporting in NPG publications that was not seen in matched manuscripts from other publishers, and this improvement appears to be ongoing. However, there is still substantial room for improvement, and this suggests that measures such as mandatory author checklists need to be supplemented by other approaches.

## Authorship: the NPQIP consortium

Study steering committee: Malcolm Macleod (University of Edinburgh, Chief Investigator and Chair), Emily

Sena (University of Edinburgh), David Howells (University of Tasmania).

Study management committee: Malcolm Macleod (University of Edinburgh, Chief Investigator and Chair), Emily Sena (University of Edinburgh), David Howells (University of Tasmania), Veronique Kiermer (Nature, until mid 2015), Sowmya Swaminathan (Nature, from mid 2015).

Redaction and identification of publications: Hugh Ash, Rosie Moreland (Imperial College, London)

Authoring and testing of training materials: Cadi Irvine, Paula Grill, Monica Dingwall, Emily Sena, Gillian Currie, Malcolm Macleod (University of Edinburgh)

Programming and data management: Jing Liao, Chris Sena (University of Edinburgh)

Outcome assessors: Paula Grill (272), Monica Dingwall (258), Malcolm Macleod (229), Cadi Irvine (179), Cilene Lino de Oliveira (170), Daniel-Cosmin Marcu (113), Fala Cramond (96), Sulail Rajani (93), Andrew Ying (81), Hanna Vesterinen (31), Roncon Paolo (28), Kaitlyn Hair (26), Marie Soukupova (23), Devon C. Crawford (17), Kimberley Wever (16), Mahajabeen Khatib (16), Ana Antonic (13), Thomas Ottavi (13), Xenios Milidonis (12), Klara Zsofia Gerlei (10), Thomas Barrett (10), Ye Liu (10), Chris Choi (9), Evandro Araújo De-Souza (8), Alexandra Bannach-Brown (8), Peter-Paul Zwetsloot (5), Kasper Jacobsen Kyng (5), Sarah McCann (4), Emily Wheater (4), Aaron Lawson McLean (1), Marco Casscella (1), Alice Carter (1), Privjyot Jheeta (1), Emma Eaton (1).

Reconciliation: Alexandra Bannach-Brown (199), Malcolm Macleod (197), Monica Dingwall (167), Paula Grill (161), Kaitlyn Hair (97), Cilene Lino de Oliveira (40), Sulail Rajani (9), Daniel-Cosmin Marcu (8), Cadi Irvine (3), Fala Cramond (1).

Data analysis: Paula Grill, Jing Liao, Malcolm Macleod

Writing Committee: Malcolm Macleod, David Howells, Jing Liao, Paul Grill, Emily Sena

Disclaimer: the opinions expressed in this article are the authors' own and do not reflect the view of any employing agency including the U.S. National Institutes of Health, the U.S. Department of Health and Human Services, or the United States Government.”

## Role of Nature in data analysis and data ownership

The study dataset belongs to the investigators, and all decisions relating to data analysis and publication were be taken by the steering committee and were independent of Nature. NPG were invited to correct any errors of fact in a draft version of the manuscript.

